# Phenotypic variability correlates with clinical outcome in *Cryptococcus* isolates obtained from Botswanan HIV/AIDS patients

**DOI:** 10.1101/418897

**Authors:** Kenya E. Fernandes, Adam Brockway, Miriam Haverkamp, Christina A. Cuomo, Floris van Ogtrop, John R. Perfect, Dee A. Carter

## Abstract

Pathogenic species of *Cryptococcus* cause hundreds of thousands of deaths annually. Considerable phenotypic variation is exhibited during infection, including increased capsule size, capsule shedding, giant cells (≥ 15 μm) and micro cells (≤ 1 μm). We examined 70 clinical isolates of *Cryptococcus neoformans* and *Cryptococcus tetragattii* from HIV/AIDS patients in Botswana to determine if the capacity to produce morphological variants was associated with clinical parameters. Isolates were cultured under conditions designed to simulate *in vivo* stresses. Substantial variation was seen across morphological and clinical data. Giant cells were more common in *C. tetragattii,* while micro cells and shed capsule occurred in *C. neoformans* only. Phenotypic variables fell into two groups associated with differing symptoms. The production of “large” phenotypes (greater cell and capsule size and giant cells) was associated with higher CD4 count and was negatively correlated with intracranial pressure indicators, suggesting these are induced in early-stage infection. “Small” phenotypes (micro cells and shed capsule) were associated with lower CD4 counts, negatively correlated with meningeal inflammation indicators and positively correlated with intracranial pressure indicators, suggesting they are produced later during infection and may contribute to immune suppression and promote proliferation and dissemination. These trends persisted at the species level, indicating that they were not driven by association with particular *Cryptococcus* species. Isolates possessing giant cells, micro cells, and shed capsule were rare, but strikingly were associated with patient death (p=0.0165). Our data indicate that pleomorphism is an important driver in *Cryptococcus* infection.

**Importance:** Cryptococcosis results in hundreds of thousands of deaths annually, predominantly in sub-Saharan Africa. *Cryptococcus* is an encapsulated yeast, and during infection cells have the capacity for substantial morphological changes, including capsule enlargement and shedding, and variations in cell shape and size. In this study we examined 70 *Cryptococcus* isolates causing meningitis in HIV/AIDS patients in Botswana in order to look for associations between phenotypic variation and clinical symptoms. Four variant phenotypes were seen across strains: giant cells ≥ 15 μm, micro cells ≤ 1 μm, shed extracellular capsule, and irregularly shaped cells. We found “large” and “small” phenotypes were associated with differing disease symptoms, indicating that their production may be important during the disease process. Overall, our study indicates that *Cryptococcus* strains that can switch on cell types under different situations may be more able to sustain infection and resist the host response.

## Introduction

Cryptococcosis, caused by pathogenic *Cryptococcus* species, is currently ranked as one of the three most common life-threatening opportunistic infections in individuals with HIV/AIDS worldwide (1, 2). The health burden is particularly high in the developing world and it is difficult to resolve despite current best antifungal therapy (2, 3).. Of annual cryptococcal-related deaths, 75% occur in sub-Saharan Africa where cryptococcal disease presents in 15-30% of HIV/AIDS patients and is associated with ∼70% mortality at 3 months (4, 5). *C. neoformans* is the major cause of disease in immunocompromised individuals, while species in the *C. gattii* complex tend to infect immunocompetent individuals (6). However, *C. gattii* complex species are increasingly being identified in HIV/AIDS patients, particularly *C. bacillisporus* and *C. tetragattii*, with the latter found associated with HIV/AIDS infections in Africa (7-10).

*Cryptococcus* yeast cells generally possess a thick capsule that is considered the major virulence factor, although this can be thin or even absent in clinical samples (11). The capsule protects the cell from phagocytosis and from reactive oxygen species damage (12, 13). Shed extracellular capsule is thought to impair the host immune response, leading to macrophage dysfunction and cell death (14). Capsule size varies greatly among *Cryptococcus* strains, and dramatically increases in response to environmental stresses including host infection (15). Cryptococcal cells can also change in size during infection (16) and various studies have emphasised the plasticity of the cryptococcal genome (17-20). Individual strains can give rise to variant populations including giant cells with cell bodies larger than 15 μm, and micro cells with cell bodies smaller than 1 μm in diameter (21, 22). These phenotypes are frequently observed *in vivo* and are likely to be important in human infection (21, 23-25).

Capsule and cell size, and the production of variants, can be experimentally modulated *in vitro* by simulating host-specific conditions, and differ between species (26). Here, we examine *in vitro* capsule and cell size variation in a collection of clinical isolates taken from HIV patients in Botswana, comprising 53 *C. neoformans* isolates across four molecular genotypes (VNI, VNII, VNBI, VNBII), 16 *C. tetragattii* (VGIV) isolates, and a single *C. gattii* (VGI) isolate. We present significant correlations between species, phenotype, and clinical outcome, explore phenotypic differences between *C. neoformans* and *C. tetragattii* that might reflect their differing pathogenesis, and show that the capacity for variation may be associated with higher virulence.

## Results

### Botswanan clinical isolates have high levels of genetic diversity

MLST analysis divided the 70 *Cryptococcus* isolates into different major species and genotypes. The collection comprised 53 *C. neoformans* isolates including genotypes VNI (n=17; 24.3%), VNII (n=2; 2.9%), VNBI (n=25; 35.7%), and VNBII (n=9; 12.9%), 17 *C. gattii* species complex isolates including *C. gattii* (n=1; 1.4%), and *C. tetragattii* (n=16; 22.9%). For simplicity we have excluded the single *C. gattii* isolate from statistical analysis. Across the collection there were 16 CAP59 ATs, 12 GPD1 ATs, 16 IGS1 ATs, 16 LAC1 ATs, 14 PLB1 ATs, 21 SOD1 ATs, and 15 URA5 ATs (Supplementary Table 1). In VNII and VNBI each isolate was assigned a unique ST, while there were 11 STs in VNI (n=16), 24 STs in VNBI (n=25), and 5 STs in VGIV (n=16) (Table 1).

**Table 1:**
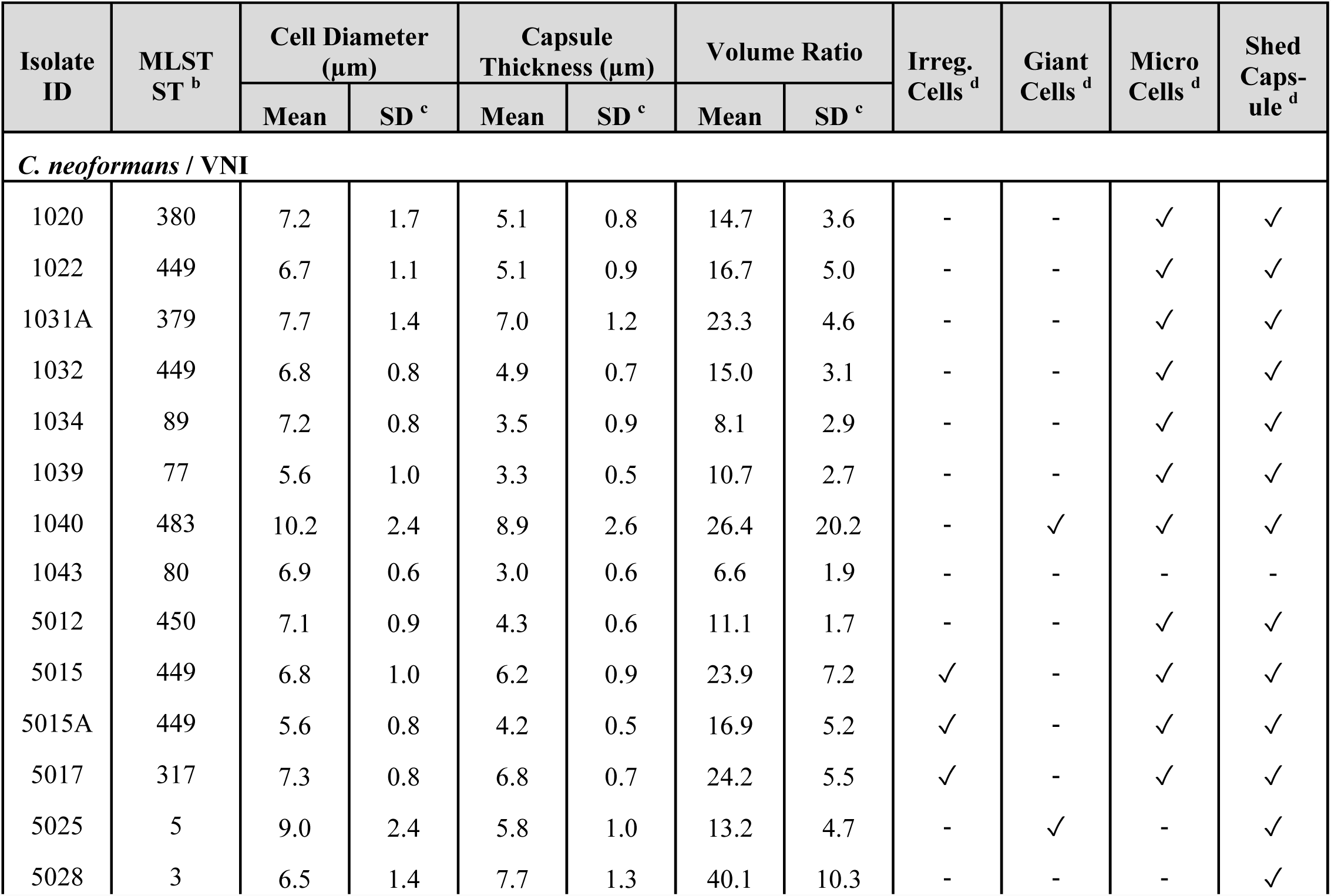
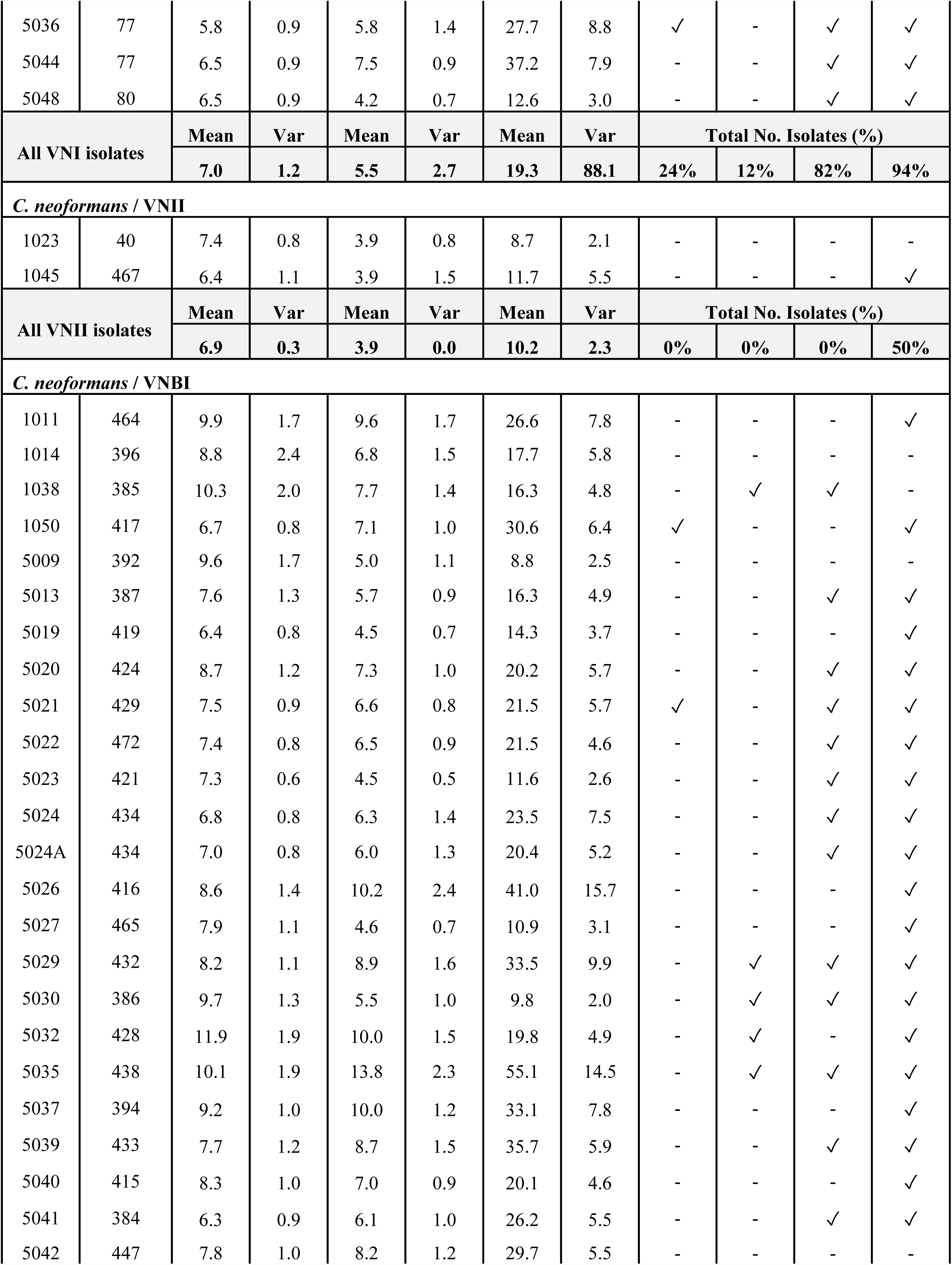
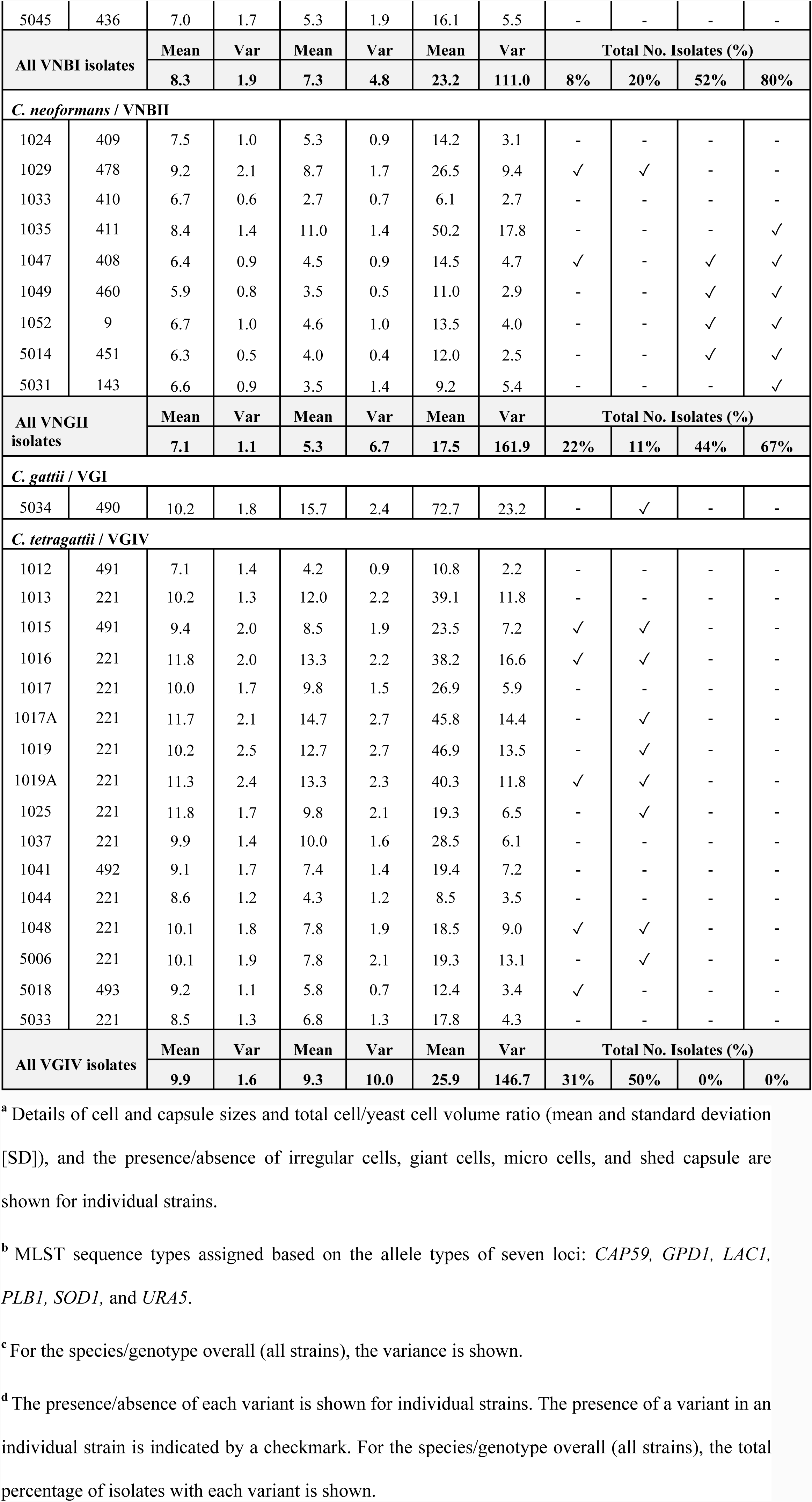
Isolates used in this study with cell and capsule size details ^a^

In order to visualise the amount of genetic variation within each genotype/species, the concatenated sequences of these seven loci were used to generate minimum spanning networks in TCS (Fig. 1). Substantial differences in genetic diversity were seen among genotypes. The *C. tetragattii*/VGIV population had a largely clonal structure with 12 out of 16 isolates belonging to ST 221 and limited divergence of the remaining isolates, while all *C. neoformans* genotypes showed much more diversity, with VNBI having the highest level. Evidence of recombination, shown as loops in the networks, was seen in the VNBI and VNBII populations.

**Figure 1:**
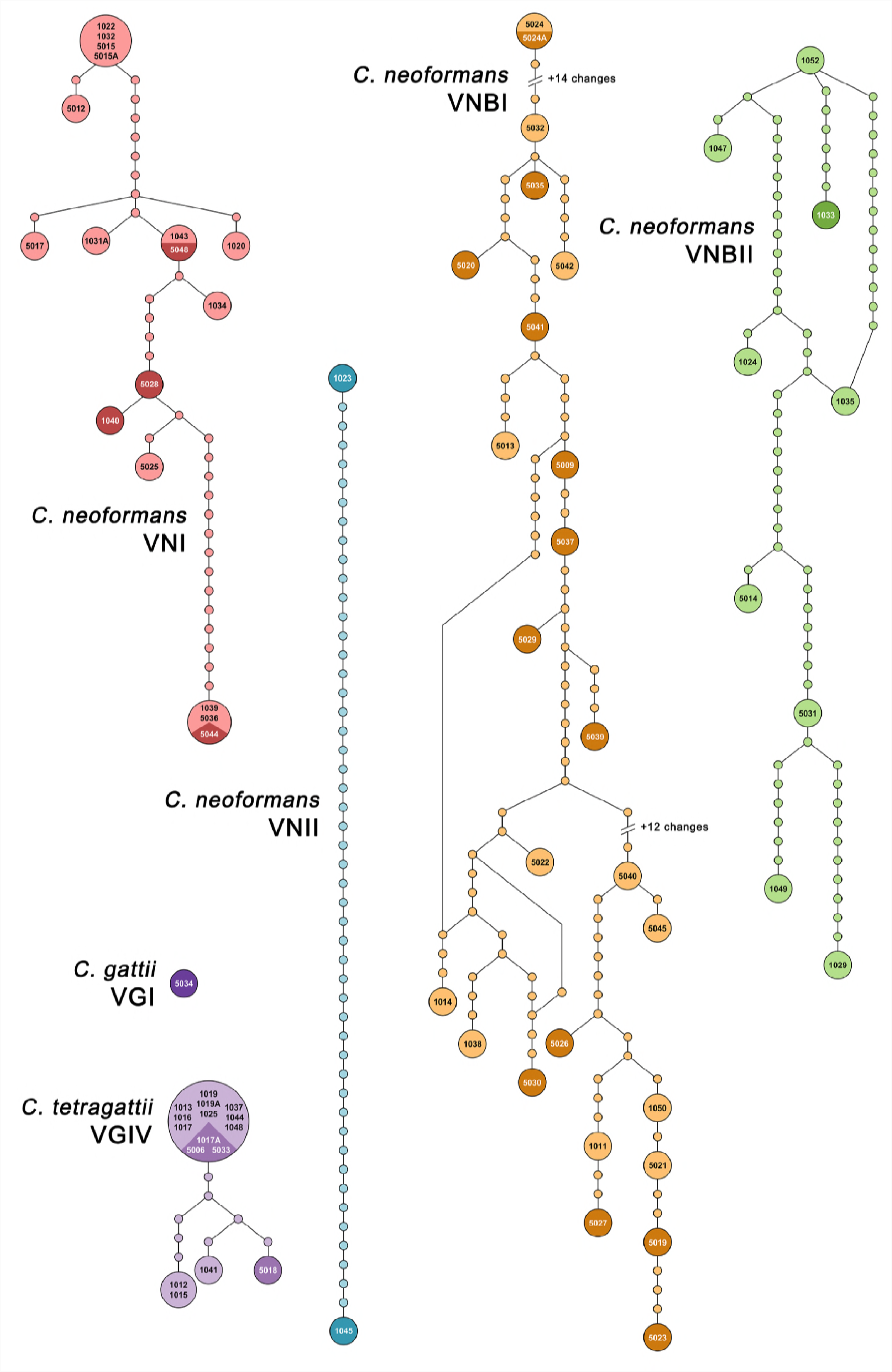
Genetic diversity and relatedness among clinical isolates belonging to each genotype. Minimum spanning network analysis of the clinical isolates based on the concatenated sequences of seven MLST loci. Each unique sequence type is represented by a circle, with size proportional to the number of isolates belonging to that sequence type. Isolate names in black type correspond to patients who were alive at the time of the analysis, while isolate names in white type on darker backgrounds correspond to patients who died. *C. neoformans* VNI n = 17, VNII n = 2, VNBI n = 25, VNBII n = 9, *C. gattii* VGI n = 1, *C. tetragatiii* VGIV n = 16. MLST alleles for each strain can be found in Supplementary Table S1.

### Induced capsule and cell size differ between species and genotypes

Capsule thickness and yeast cell diameter were measured after growth in DMEM broth with 5% CO_2_ at 37 ^°^C for 5 days, and data were compared across genotypes for mean and variation (Table 1; Fig. 2A-C). On average, *C. tetragattii* isolates had significantly greater capsule thickness (p=0.0025) and yeast cell diameter (p<0.0001) than *C. neoformans* isolates. Within *C. neoformans* genotypes, VNBI isolates had significantly greater capsule thickness than VNI isolates (p=0.0043) and significantly greater yeast cell diameter than VNI (p=0.0032) and VNBII (p=0.0178) isolates. F test analysis was used to compare the variance of the data (Table 1). Despite their limited genetic diversity, *C. tetragattii* isolates had significantly more variation in capsule thickness than *C. neoformans* isolates (p=0.0245). There were no significant differences in variation in capsule thickness among *C. neoformans* genotypes. Yeast cell diameter measurements did not vary significantly different between any groups.

**Figure 2:**
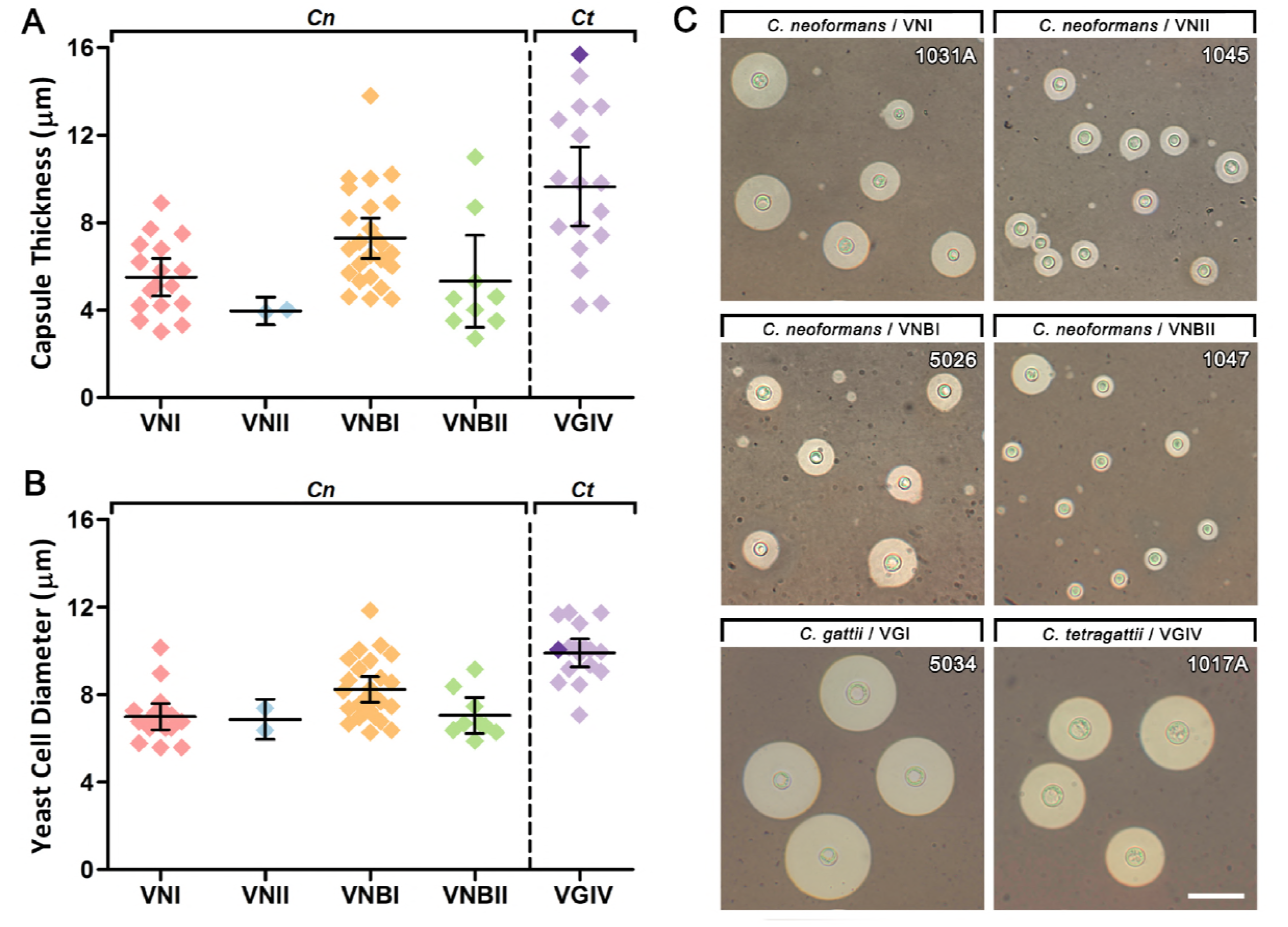
Capsule thickness and yeast cell diameter vary among the clinical isolates following growth under capsule-inducing conditions. **(A)** Capsule thickness and **(B)** yeast cell diameter of clinical isolates from each genotype grown on DMEM with 5% CO_2_ at 37 ^°^C for 5 days. Each data point represents the average of 100 cells measured for a single isolate. Error bars show the mean ± 95% confidence interval. *C. neoformans* VNI n = 17, VNII n = 2, VNBI n = 25, VNBII n = 9, *C. gattii* VGI n = 1 (dark purple), *C. tetragattii* VGIV n = 16 (light purple). **(C)** Indian ink preparations of representative strains from each genotype showing variation in capsule and cell size. Scale bar = 30 μm.

### Giant cells are more frequent in *C. tetragattii* while micro cells and extracellular capsule are present only in *C. neoformans*

Following growth under inducing conditions, the three variant phenotypes seen at differing frequencies across isolates were giant cells (yeast cell diameter ≥ 15 μm) (Fig. 3A), micro cells (yeast cell diameter ≤ 1 μm) (Fig. 3B) and shed capsule (Fig. 3C). Elongated and irregularly shaped cells were seen in a number of strains across all genotypes (except VNII where n=2) (Supplementary Figure S1D, Table 1) but these comprised a small subset (<5%) of the total population of cells. Cells larger than 15 μm have previously been classed as “titan cells”, however as *in vivo* titan cells possess additional defining characteristics that were not measured in this study including altered capsular structure, increased DNA content and increased vacuolar formation (27), they will be here referred to as giant cells. The number of clinical isolates exhibiting these morphological variants for each genotype is summarised in Table 1. Giant cells were significantly associated with greater capsule thickness (p<0.0001) and yeast cell diameter (p<0.0001) indicating that isolates with larger cells are more likely to produce giant cells, and Fig. 2B shows that cell size is generally spread along a continuum. Micro cells and shed capsule, however, were not significantly associated with capsule thickness or yeast cell diameter in *C. neoformans* indicating that cell size is not related to their production and that micro cells are a distinct cell type rather than the endpoint of a continuum of increasingly smaller cells.

**Figure 3:**
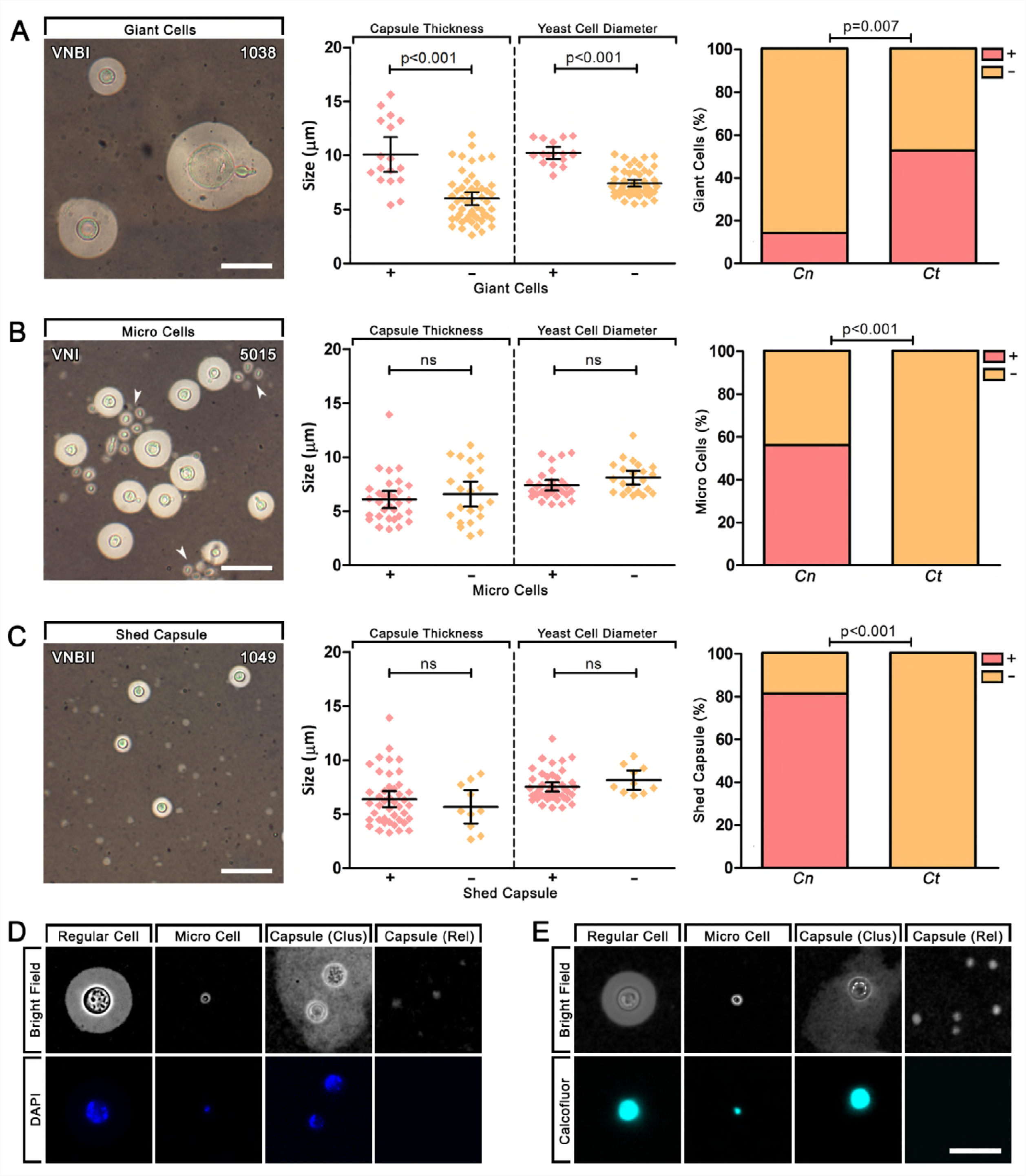
Morphological variants produced by clinical isolates following growth under capsule-inducing conditions. Indian ink preparations, capsule and cell size associations, and species associations of morphologically variant phenotypes including **(A)** giant cells > 15 μm, **(B)** micro cells < 1 μm, and **(C)** shed capsule. Scale bars = 20 μm. Error bars show the mean ± 95% confidence interval. Indian ink preparations were further stained with **(D)** DAPI or **(E)** Calcofluor White (CFW) to examine nuclei and cell walls respectively. In isolates that were scored as having shed capsule, this was occasionally present as an irregular cluster around cells (Clus) as well as released into the media in small clumps (Rel). Scale bars = 15 μm.

The production of giant cells was significantly associated with *C. tetragattii* (50% of isolates; p=0.0070) compared to *C. neoformans* (15% of isolates) while micro cells and shed capsule were only seen in *C. neoformans* (58 and 81% of isolates, respectively) and were significantly associated with each other (p<0.0001). Within *C. neoformans*, giant cells, micro cells, and shed capsule were observed in all genotypes (except VNII, which is inconclusive as n=2). To confirm that micro cells and shed capsule are two distinct phenotypes, isolates were stained with DAPI (Fig. 3D) and calcofluor white (Fig. 3E) post capsule induction to investigate nuclear and cell wall morphology, respectively. Micro cells displayed fluorescence comparable to regular cells, demonstrating that they have nucleic acid material and intact cell walls. Shed capsule clustered around the cell or released into the medium displayed no fluorescence.

### Correlations occur among clinical, phenotypic, and genotypic variables

Correlations were assessed between clinical, phenotypic, and genotypic variables across the isolate collection. Correlation plots were generated showing the direction, strength, and statistical significance of correlations between clinical variables overall (Fig. 4A), between phenotypic variables overall (Fig. 4B), and between clinical and phenotypic variables overall and, for certain clinical variables, for *C. neoformans* and *C. tetragattii* species individually (Fig. 4C). All other analyses and individual p-values are recorded in Supplementary Table 2. Principal component analysis (PCA) was used to investigate the variance and dimensionality of the dataset. PCA biplots were generated using the first two components for clinical data accounting for 25.4% of variation (Fig. 4D) and for phenotypic data accounting for 61.9% of variation (Fig. 4E).

**Figure 4:**
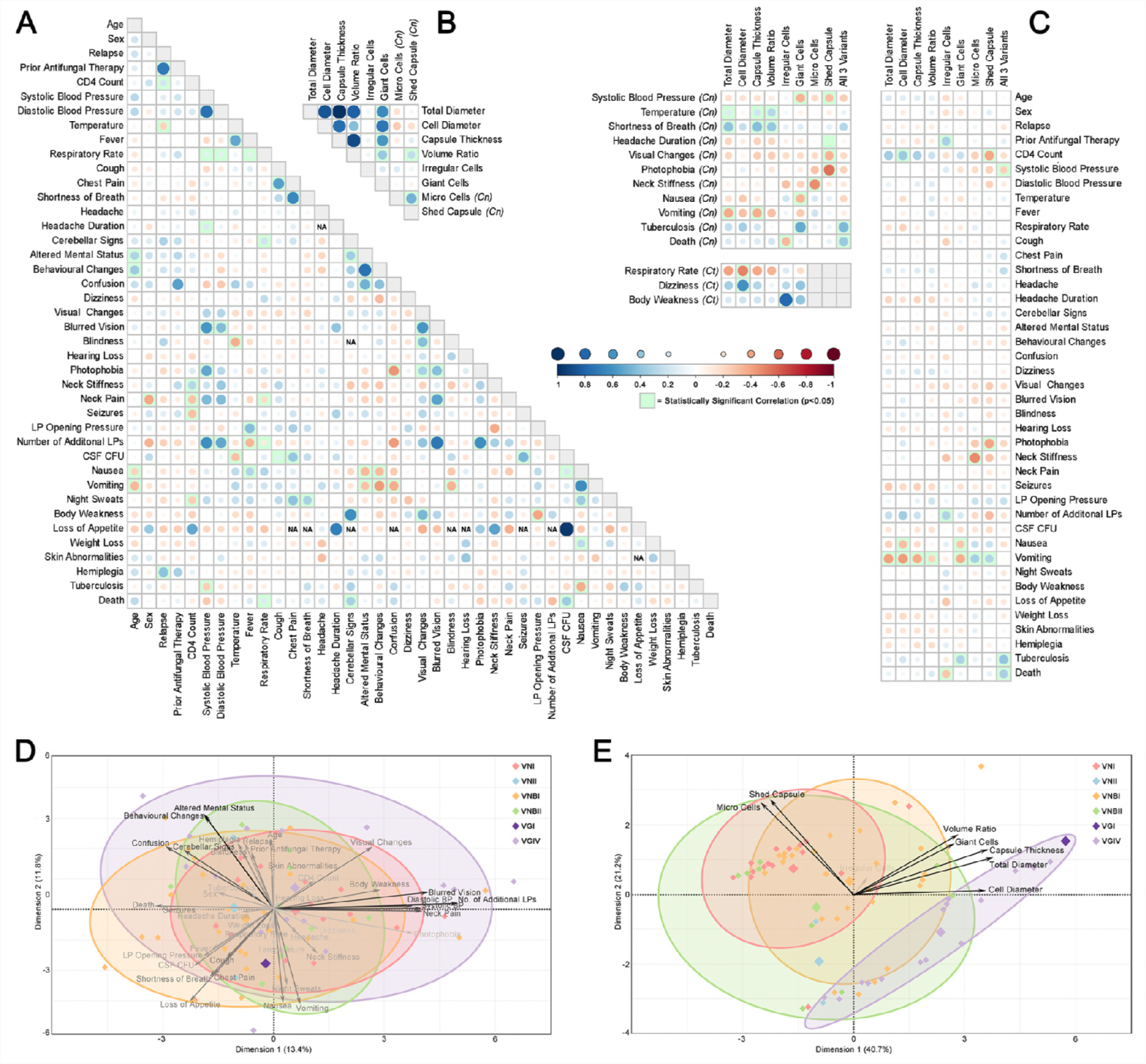
Correlation analysis of clinical and phenotypic variables. Correlation plots showing the strength and direction of associations **(A)** between clinical variables, **(B)** between phenotypic variables, and **(C)** between clinical and phenotypic variables for all isolates in the dataset. The size of the circle corresponds to the strength of the association and statistically significant associations (p<0.05) are highlighted in green. P-values for each correlation including species breakdowns can be found in Supplementary Table S2. PCA biplots of the first two significant dimensions obtained using principal component analysis of **(D)** clinical data accounting for 25.4% of variation in the dataset, and **(E)** phenotypic data accounting for 61.9% of variation in the dataset. Large diamonds represent genotype averages, length and opacity of arrows represents the degree of contribution of that variable to the model, and genotype ellipses represent the 80% confidence interval.

Most of the significant associations found between clinical symptoms were as expected, with similar variables being positively correlated including fever and temperature (p<0.0001), cough and chest pain (p=0.0013), altered mental status and behavioural changes (p<0.0001), and nausea and vomiting (p<0.0001). Similarly, cognitive symptoms such as confusion, behavioural changes, and altered mental status; respiratory symptoms such as cough, shortness of breath, chest pain, and respiratory rate; and intracranial symptoms such as visual changes, blurred vision, neck pain, and photophobia grouped closely together in the PCA biplot of clinical variables (Fig 4D). Given the complexity of clinical data that relies on self-reporting by patients that may be quite ill and subsequent interpretation by clinicians, this serves as a as a useful internal control for data quality. Patient death was significantly positively correlated with respiratory rate (p=0.0101), cerebellar signs (p=0.0478) and CSF CFU (p=0.0048).

### The capacity for production of “large” and “small” phenotypes correlates with certain clinical symptoms indicative of early and late stage infection, while the capacity for variation is associated with patient death

In the PCA biplot of phenotypic variables (Fig. 4E), properties based on cell and capsule size grouped closely together along with giant cells, while micro cells and shed capsule grouped closely with each other and away from the other properties. These groups were a strong driver of variation in the biplot, indicating genotype-specific differences, while the production of irregular cells did not. When correlated with clinical data, the phenotypic variables within these groups showed similar directions and strengths of association, with “large” phenotypes including greater capsule thickness, greater yeast cell diameter, and giant cells generally grouping and often opposing the “small” phenotypes of micro cells and shed capsule (Fig. 4C).

CD4 T-cell count is a reliable predictor of host immune status, with counts <500 indicating immune suppression, and <200 in HIV-infected individuals indicating AIDS. In the current study, CD4 counts ranged from 2-389 (Supplementary Table 1). Overall, CD4 count was positively correlated with cell size across the entire dataset (p=0.0250) and negatively correlated with the production of shed capsule (p=0.0158). Nausea and vomiting are frequent symptoms of increased intracranial pressure during cryptococcosis and these were negatively correlated with “large” cell phenotypes: nausea was negatively correlated with yeast cell diameter (p=0.0135) and giant cells (p=0.0357), and vomiting was negatively correlated with all “large” phenotypes including total diameter (p=0.0007), yeast cell diameter (p=0.0010), capsule thickness (p=0.0014), volume ratio (p=0.0269) and giant cells (p=0.0041). Vomiting was also positively correlated with the “small” phenotypes including micro cells (p=0.0077) and shed capsule (p=0.0153). Shed capsule was also negatively correlated with visual changes (p=0.0358) and photophobia (p=0.0032), symptoms typically attributed to meningeal irritation. A similar pattern was seen with fever and neck stiffness, which are associated with an aggressive inflammatory response. Patient temperature was positively correlated with total diameter (p=0.0412), capsule thickness (p=0.0104) and volume ratio (p=0.0016) in the *C. neoformans* isolates, while neck stiffness correlated negatively with micro cells across the entire collection (p=0.0041).

Finally, patient death was significantly positively correlated with the production of all three major morphological variants (p=0.0165); four patients had isolates that produced giant cells, micro cells and shed capsule, and all four patients died during the period when clinical data were scored. The production of all three major morphological variants was also positively correlated with the patient having tuberculosis (p=0.0133), and negatively correlated with systolic blood pressure (p=0.0474). In contrast, irregular cells were negatively correlated with death (p=0.0241). These were also more likely to be produced by isolates obtained from patients who had undergone antifungal therapy with either fluconazole or amphotericin B prior to admission (p=0.0438). All of the above associations were tested for each species alone as well as across the entire dataset, and the trends remained largely the same, although some statistical power was lost due to smaller sample sizes, indicating that these results were not being driven by an association with one of the species.

### Some correlations show species-specificity

Several species-specific correlations were found, indicating that aspects of pathogenesis and host response are distinct between *C. neoformans* and *C. tetragattii* isolates. In the PCA biplot of phenotypic variables, the *C. tetragattii* isolates grouped mostly separately from *C. neoformans* isolates due to their larger cell and capsule sizes, higher incidence of giant cells, and lack of micro cells and shed capsule (Fig. 4E). *C. neoformans* isolates were significantly associated with lower CD4 count (p=0.0103), higher LP opening pressure (p=0.0208) and vomiting (p=0.0013) compared to *C. tetragattii*. Isolates from the *C. neoformans* genotype VNBI were significantly associated with lower CD4 count (p=0.0063) and lower diastolic blood pressure (p=0.0235) compared to all other isolates (Supplementary Table 2). In the *C. neoformans* population, shortness of breath was positively correlated with total diameter (p=0.0430), capsule thickness (p=0.0366) and volume ratio (p=0.0367). In *C. tetragattii* isolates, yeast cell diameter was positively correlated with dizziness (p=0.0292) and negatively correlated with respiratory rate (p=0.0335).

## Discussion

Our study aimed to correlate phenotypic variation in cryptococcal isolates with pathogenesis and clinical manifestation of disease using *in vitro* stresses designed to simulate those encountered during human infection. The observed phenotypes were a stable attribute of the isolates, although we cannot be certain that these phenotypes would occur in infected patients in the same way. Furthermore, the clinical dataset used in this study contains missing values, and certain parameters rely on self-reporting, which may not be very robust. Despite this, we saw some strong correlations and trends, and while this does not necessarily imply causation, the strength and pattern of the associations indicates that phenotypic plasticity and morphological presentation do play a role in cryptococcal disease.

### Phenotypic plasticity is high and not related to genetic diversity

Almost a quarter of the clinical isolates in this study were identified as *C. tetragattii,* a species which is uncommon globally but has been found to cause a relatively high incidence of disease in HIV/AIDS patients in sub-Saharan Africa, having been identified in Botswana, Malawi, South Africa, and Zimbabwe (10, 28-30). Relatively little is known about this species, however our previous study of the *C. gattii* complex found that *C. tetragattii* is similar to other species in the complex in terms of capsule production and cell size, but along with *C. bacillisporus* is more temperature sensitive than virulent genotypes *C. gattii* and *C. deuterogattii*, and commonly produces irregular cells. This suggests that *C. tetragattii* is a “weaker” pathogen and may be infecting immunocompromised hosts due to high environmental presence in Botswana and other sub-Saharan African countries. Environmental studies of the area are needed to determine where its niche lies.

MLST analysis revealed a limited amount of genetic diversity among *C. tetragattii* isolates, which contrasted with high genetic diversity and evidence of recombination in the *C. neoformans* genotypes (Table 1; Fig. 1), something that has been seen in other studies of southern African *C. neoformans* populations (29, 31-33). Significant phenotypic differences occurred among, between and within the *C. neoformans* genotypes and within *C. tetragattii* and, despite their clonal nature, *C. tetragattii* isolates had significantly more variation in capsule thickness (p=0.0245), suggesting that some phenotypic differences may be a result of epigenetic mechanisms. Rhodes et al. (2017) found the most rapidly evolving genes between lineages of *C. neoformans* to be transcription factors and transferases, suggesting that transcriptional reprogramming and remodelling may be responsible for generating phenotypic diversity, rather than genomic changes (34). Phenotypic plasticity allows rapid adaptation inside the host and is common in many fungal species, such as *Candida albicans*, which can exhibit various morphological types including yeast and filamentous forms, and white and opaque forms which influence mating, virulence, and interactions with immune cells *in vitro* (35). *Cryptococcus*, like *Candida*, appears to show a level of pleomorphism with different phenotypes appearing in response to stress.

### Clinical markers of early and late infection suggest cell phenotypes change during the course of infection and may play a role in immune response

A model of the relationship between the various phenotypic variants, cryptococcal species, and clinical variables based on the significant associations and trends found in this study is presented in Fig. 5. CD4 count, representing the immunity status of the patient, was overall positively correlated with “large” phenotypes suggesting that larger cells, more capsule, and giant cells are produced during earlier stages of infection, and overall negatively correlated with “small” phenotypes suggesting that micro cells and shed capsule may be a later response during infection as immune function declines (Fig. 4C; Fig. 5). Nausea and vomiting are symptoms that are typically associated with increased intracranial pressure as this results in stimulation of the vomiting center of the brain. These were overall negatively correlated with “large” phenotypes again suggesting that larger cells may be an earlier response, and overall positively correlated with “small” phenotypes. The latter could be expected as shed capsule in particular has long been implicated in increasing intracranial pressure due to the accumulation of capsular polysaccharide blocking passage of the CSF across arachnoid villi (36). In all, this suggests a transition from “large” to “small” cell phenotypes as infection progresses and intracranial pressure rises.

**Figure 5:**
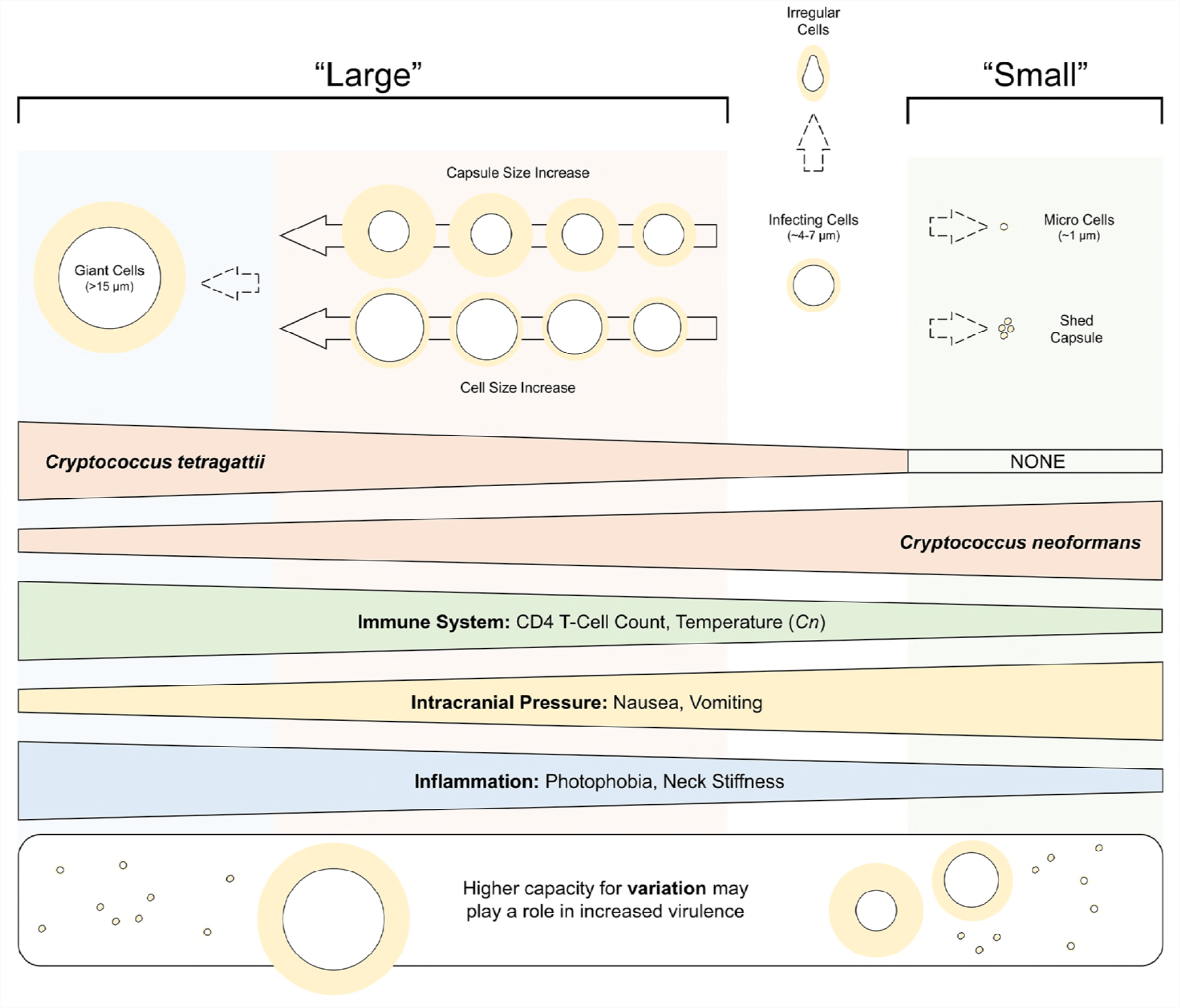
A summary model of the phenotypic variants seen in this study and the direction of their associations with the *C. tetragattii*, *C. neoformans*, and clinical variables. “Large” phenotypes include larger yeast cell size, larger capsule size, and giant cells; these are prevalent in *C. tetragattii* and are generally correlated with symptoms indicating a less suppressed immune system and low intracranial pressure. “Small” phenotypes include micro cells and shed capsule; these are prevalent in *C. neoformans* and are generally correlated with symptoms indicating a more suppressed immune system, high intracranial pressure, and low inflammation. Overall, a higher capacity for variation may play a role in increased virulence in *Cryptococcus.*

The “small” phenotypes were also overall negatively correlated with visual changes (significantly with shed capsule, p=0.0358) and photophobia (significantly with shed capsule, p=0.0032), symptoms associated with inflammation of the meninges (43). It has been observed that cryptococcal meningitis in HIV patients is characterised by a lack of an active host inflammatory response (36) so it is possible that micro cells and extracellular capsule accumulating in the CSF and causing raised intracranial pressure might also be dampening the host inflammatory response. The major cryptococcal capsular polysaccharide (GXM), has been documented to have numerous immunosuppressive properties including modulation of cytokine secretion, induction of macrophage apoptosis, and suppression of leukocyte migration (37-40). Studies in mice have found GXM to markedly dampen the hyperinflammatory response via inhibition of proinflammatory cytokine secretion (41), and the administration of purified GXM reduced the number of immune cells infiltrating the brain (42)

Capsule size in cryptococcal cells was found to differ significantly between genotypes, species, and individual isolates (Table 1; Fig. 2A-C). Compared to all *C. neoformans* genotypes, *C. tetragattii* isolates had significantly thicker capsules (p=0.0025) and larger yeast cells (p<0.0001). Studies to date investigating the relationship between capsule size and virulence have found conflicting results. Robertson et al. (2014) found highly encapsulated *C. neoformans* isolates were significantly associated with lower fungal clearance rates, poor inflammatory responses, and increased intracranial pressure (43). In contrast, Pool et al. (2013) showed that hypercapsular *C. neoformans* isolates possessed the least neurovirulence in a mouse model, while isolates producing less capsule were more virulent and resulted in a higher fungal load in the brain (44). As such, it is unclear whether a large capsular phenotype enhances the overall virulence of *Cryptococcus*. Our current study found that capsule size strongly correlates with yeast cell size, and it is possible that capsule alone has less importance than the overall size of the cell.

### Giant cells are prevalent in *C. tetragattii* and likely result from a gradual increase in cell size while micro cells appear exclusive to *C. neoformans* and are a distinct cell population strongly associated with shed capsule

The presence of giant cells has been recorded in infected tissues during mammalian infection, where they appear most frequently in the extracellular space (21, 22, 25, 45). Giant cells are thought to be important in the establishment and persistence of infection due to their large size preventing phagocytosis and allowing them to remain in the host for long periods of time (16). Giant cells also appear to increase virulence through conferring resistant properties to their normal size progeny enhancing their capacity for survival and dissemination and thus increasing the overall virulence of the strain (16, 45). The vast morphological changes associated with *in vivo* giant cell production, including alteration of cell body and organelles, indicate that giant cell production is a developmental transition (27, 46). Recent studies on giant cell production found that the capacity to produce giant cells *in vitro* may be a reliable predictor of their formation *in vivo* (47), and that regularly sized cells present in the initial inoculum transitioned progressively towards the giant cell phenotype under inducing conditions (48) with the transition occurring at low cell densities (49). The current study found giant cells to be significantly associated with larger average yeast cell diameters (p<0.0001) and greater capsule thickness (p<0.0001) in both *C. neoformans* and *C. tetragattii* (Fig. 3A) which supports the hypothesis of a gradual shift towards a larger phenotype. Although the majority of literature on giant cells relates to *C. neoformans*, we found that the presence of giant cells was significantly associated with the *C. tetragattii* complex.

*Cryptococcus* micro cells are an intriguing and understudied class of cell that has to date received little attention. These cells are commonly seen in infection and have been speculated to assist in the infection process, with their small size allowing them cross biological barriers and to disseminate easily to the brain (14, 44). The presence of micro cells was strongly (p<0.0001), but not always, associated with shed capsule (Table 1; Supplementary Table 2) suggesting that they are induced by the same or similar processes. Staining with DAPI and calcofluor white, confirmed that micro cells are real cells with nuclear material and cell walls, and are distinct from shed capsule (Fig. 3D-E). Furthermore, unlike giant cells, micro cells appear to be a distinct cell class as there was no correlation with cell size, indicating that there is no continuum of increasingly smaller cells. Their presence appears to be exclusive to *C. neoformans* (and seen across genotypes); the current analysis and our previous study investigating 70 *C. gattii* complex strains including *C.* g*attii*, *C. deuterogattii*, *C. bacillisporus*, and *C. tetragattii* clinical, environmental, and veterinary strains and found no evidence of micro cells in any strain (26).

### Pleomorphism may have a role in the virulence of *Cryptococcus*

The production of “large” and “small” phenotypes was largely mutually exclusive; most isolates possessed either giant cells or micro cells/shed capsule but rarely both. It is therefore intriguing that the four isolates that produced all three major morphological size variants (giant cells, micro cells, and shed capsule) resulted in patient death (p=0.0165) (Fig. 4C). While this must be viewed with the limitation that there were very few isolates within this category, it may indicate that the capacity for variation plays a role in virulence. Three out of four of these isolates belonged to VNBI, which appeared to be the most virulent genotype with the highest percentage of patient deaths, and also with the largest average capsule sizes within *C. neoformans*. As “large” and “small” variants appear associated with quite different disease symptoms, this could enable greater capacity for infection, immune evasion, and pathogenesis. An alternative hypothesis is that in the severely weakened immune state of late HIV/AIDS patients, different cryptococcal cell types can flourish; however many isolates caused patient death but did not demonstrate this diversity of cell types. Further work into the factors that induce each of these unique cell types and their role in virulence and disease progression is required to substantiate this preliminary finding.

Conversely, the presence of “irregular cells” with irregularly shaped and elongated cell morphologies was significantly negatively correlated with death (p=0.0241) suggesting that these are defective cells. Irregular cells were not associated with any particular species or genotype and had no significant correlations with any other phenotypic variable, but they were significantly more likely to occur in isolates isolated from patients that had undergone antifungal therapy prior to admission (p=0.0438). This suggests that irregular cells may be produced by isolates with reduced drug susceptibility but with a cost of lower resistance to host-imposed stress, making them less able to mount aggressive disease. *In vitro*, we have found the production of these cells may be triggered by the stress of nutrient limitation (26). The extent of the elongation of these cells varied from isolate to isolate, but in some cases they appeared to approach the morphology of pseudohyphal forms. Pseudohyphal forms have been reported in *Cryptococcus* but are thought to be rare during cryptococcosis and are also associated with reduced virulence (27, 50).

This study demonstrates the complex relationship between phenotypic variation and adaptation to the host environment in *Cryptococcus*, with pleomorphic characters potentially contributing to overall virulence. As different properties may be beneficial at different stages and sites of infection, isolates that are able to produce diverse cells in response to changing situations may be more able to sustain infection and resist the host response.

## Materials & Methods

### *Cryptococcus* isolates

A collection of 70 *Cryptococcus* isolates were provided from two major public hospitals in Botswana: the Princess Marina Hospital in Gaborone and the Nyangbwe Referral Hospital in Francistown, as part of an ongoing study into cryptococcosis in Africa. Some of this collection has been reported previously in Chen et. al (2015) (29). Fungal cells were cultured from the cerebrospinal fluid (CSF) of patients with HIV/AIDS and cryptococcal meningitis enrolled within an 18-month period from 2012-2013. Cryptococcal meningitis was confirmed by a positive India ink test or CSF culture. Patients received induction therapy with amphotericin B, however all isolates were obtained before treatment commenced. Any patients where death was not attributable to cryptococcal meningitis were excluded from the dataset. Screening on fluconazole plates found no isolates with high-level resistance. All isolates used in this study are referred to by their isolate ID and are listed in Table 1, with full details of isolate name, genotypic, phenotypic, and clinical data listed in Supplementary Table 1. Phenotypic data for type strain H99 is also included in this table as a reference.

### Culture conditions

*Cryptococcus* isolates were cultured from –80 ^°^C glycerol stocks, streaked for single colonies on Sabouraud Dextrose Agar (SDA; 10 g peptone, 40 g glucose, 15 g agar, 1 L dH_2_O) and incubated at 30 ^°^C for 48 hours. To standardise growth phase, each isolate was grown overnight from a single colony in 10 ml of Sabouraud Dextrose Broth (SDB; 10 g peptone, 20 g glucose, 1 L dH_2_O) in a 100 ml Schott bottle at 37 ^°^C with 180 rpm shaking until the culture reached exponential growth phase.

### Capsule induction

Capsule induction was optimised using various media designed to mimic stresses encountered during mammalian infection that are frequently used for *C. neoformans*, and media developed during a previous study in the *C. gattii* complex (25). This included Dulbecco’s Modified Eagle Medium (DMEM; Life Technologies), and Sabouraud medium in both plate (SDA) and broth (SDB) form diluted 10-fold (CIM-10), or 20-fold (CIM-20) in 50 mM MOPS (Sigma Aldrich) (13, 26). For culture on agar plates, a single loopful of cells was taken from the overnight culture and streaked for single colonies onto each medium. For broth cultures, cells were collected by centrifugation, washed once with Phosphate-Buffered Saline (PBS; Oxoid), and counted with a haemocytometer before 10^5^ cells were inoculated into 5 ml of media in a 6-well tissue culture plate (BD Falconer). All cultures were incubated with 5% CO_2_ at 37 ^°^C for 5 days. As a control, isolates were streaked for single colonies on SDA and incubated at 30 ^°^C for 5 days. DMEM broth was the most successful induction media (Supplementary Figure S1) and was used for all subsequent analyses.

### Staining and microscopy

To visualise capsule, single colonies from plates or 1 ml of culture from broths were suspended or resuspended in 150 μl of PBS and counterstained with 20 μl of India Ink. A 15 μl aliquot of this mixture was placed on a glass slide and dried for 10 min under a coverslip. Slides were then photographed using an IS10000 Inverted Microscope (Luminoptic) under the 40x objective using ISCapture Imaging software (Tucsen Photonics). For each isolate, a minimum of 20 random fields of view were photographed using stage coordinates determined by a random number generator. Additional stains used to visualize nucleus and cell wall morphology respectively were: (i) DAPI (Sigma Aldrich) at 1:5000 incubated for 30 minutes at room temperature, (ii) Calcofluor White (Sigma Aldrich) at 1 g/l with one drop of 10% potassium hydroxide incubated for 2 minutes at room temperature.

### Measurement of cell and capsule size

Total diameter (including capsule) (*d*_*t*_) and yeast cell diameter (*d*_*y*_) were measured for 100 cells per isolate using ImageJ software (National Institutes of Health). From these measurements, capsule thickness (*t*_*c*_) was calculated as 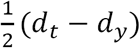. Total volume (*v*_*t*_) and yeast cell volume (*v*_*y*_) were calculated using the formula for the volume of a sphere 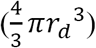. Cells with *d*_*y*_ greater than 15 μm or less than 1 μm were identified as giant cells or micro cells, respectively, and were noted along with any morphologically irregular cells. These variants were excluded from all assessments of mean cell size for isolate populations.

### MLST analysis

Genetic variation was studied using the International Society of Human and Animal Mycology (ISHAM) consensus MLST scheme for the *C. neoformans*/*C. gattii* species complex, which uses seven unlinked genetic loci: the housekeeping genes *CAP59, GPD1, LAC1, PLB1, SOD1,* and *URA5*, and the non-coding intergenic spacer region *IGS1* (51). Sequences were obtained from SNPs identified from whole genome sequences, where available (52). Remaining loci were amplified independently using ISHAM recommended primers and amplification conditions, and PCR products were commercially purified and sequenced by Macrogen Inc. (Seoul, Korea). Sequences were edited using Geneious R6 (Biomatters Ltd.). An allele type (AT) was then assigned for each of the seven loci per isolate and the resulting allelic profile used to assign a sequence type (ST) according to the ISHAM consensus MLST database (http://mlst.mycologylab.org). A minimum spanning network of the concatenated sequences was generated using the TCS 1.21 software package (http://darwin.uvigo.es/software/tcs.html) to visualise the relatedness of isolates (53).

### Statistical analysis

Significant differences between species or genotypes for phenotypic or clinical data were determined using two-tailed unpaired t-tests with Welch’s correction. Differences in variance between species and genotypes were assessed by F test analysis. Associations between continuous phenotypic and clinical variables used Spearman rank-order correlations, those between continuous and binary variables used Mann-Whitney *U* tests, and those between binary variables used chi-square tests, or Fisher’s exact tests if any expected value was <5. Correlations were tested across all isolates, across *C. neoformans* isolates only, and across *C. tetragattii* isolates only, with all p values listed in Supplementary Table 2. P values of <0.05 were considered significant. Error bars represent the mean +/-95% confidence intervals. Data were analyzed using Excel (Microsoft Corporation), Prism 5 (GraphPad Inc.), and SPSS Statistics (IBM) software. Correlation plots and principal component analysis (PCA) biplots were generated in R 3.4.0 (R Core Team) using the packages corrplot for correlation plots, missMDA to impute missing values (54), and FactoMineR and factoextra to generate PCA biplots (55).

## Author Contributions

KEF, DAC and JRP conceived and designed the experiments. KEF and AB produced the phenotype data; KEF and CAC produced the genotype data; and all clinical data was collected by MH and JRP. KEF collated and analysed the data and wrote the manuscript with assistance from DAC and JRP. FvO assisted with statistical analysis.

---

**Supplementary Figure S1: Capsule enlargement on different media and morphologically irregular cells produced following growth on DMEM. (A)** Indian ink preparations of *C. neoformans* VNI strain H99 showing capsule enlargement when grown using various induction media compared to growth on SDA at 30 ^°^C. **(B) C**apsule thickness and **(C)** yeast cell diameter of 50 individual cells from eight randomly selected clinical isolates grown in CIM-20 vs DMEM broth. Error bars show the mean ± 95% confidence interval. **(D)** Induction media DMEM induced occasional irregular, elongated cells in some strains. Scale bars = 30 μm.

**Table Headings**

**Table 1:** Isolates used in this study with cell and capsule size details ^a^

**Supplementary Table S1:** Genetic, phenotypic, and clinical data for all isolates used in this study.

**Supplementary Table S2:** Statistical significance for all associations tested in this study.

